# Herpes simplex virus 1 expressing GFP-tagged virion host shutoff (vhs) protein uncouples the activities of degradation and nuclear retention of the infected cell transcriptome

**DOI:** 10.1101/2022.01.04.475014

**Authors:** Emma L Wise, Jerzy Samolej, Gillian Elliott

**Affiliations:** Section of Virology, Department of Microbial Sciences, Faculty of Health and Medical Sciences, University of Surrey, Guildford, United Kingdom

## Abstract

Virion host shutoff (vhs) protein is an endoribonuclease encoded by herpes simplex virus 1 (HSV1). Vhs causes a number of changes to the infected cell environment that favour translation of late (L) virus proteins: cellular mRNAs are degraded, immediate-early (IE) and early (E) viral transcripts are sequestered in the nucleus with polyA binding protein (PABPC1), and dsRNA is degraded to help dampen the PKR-dependent stress response. To further our understanding of the cell biology of vhs, we constructed a virus expressing vhs tagged at its C-terminus with GFP. When first expressed, vhs-GFP localised to juxtanuclear clusters, and later it colocalised and interacted with its binding partner VP16, and was packaged into virions. Despite vhs-GFP maintaining activity when expressed in isolation, it failed to degrade mRNA or relocalise PABPC1 during infection, while viral transcript levels were similar to those seen for a vhs knockout virus. PKR phosphorylation was also enhanced in vhs-GFP infected cells, in line with a failure to degrade dsRNA. Nonetheless, mRNA FISH revealed that as in Wt but not Δvhs infection, IE and E, but not L transcripts were retained in the nucleus of vhs-GFP infected cells at late times. Moreover, a representative cellular transcript which is ordinarily highly susceptible to vhs degradation, was also retained in the nucleus. These results reveal that the vhs-induced nuclear retention of the infected cell transcriptome is dependent on vhs expression but not on its endoribonuclease activity, uncoupling these two functions of vhs.

**Importance:** Like many viruses, herpes simplex virus 1 (HSV1) expresses an endoribonuclease, the virion host shutoff (vhs) protein, which regulates the RNA environment of the infected cell and facilitates the classical cascade of virus protein translation. It does this by causing the degradation of some mRNA molecules and the nuclear retention of others. Here we describe a virus expressing vhs tagged at its C-terminus with green fluorescent protein (GFP) and show that the vhs-GFP fusion protein retains the physical properties of native vhs, but does not induce the degradation of mRNA. Nonetheless, vhs-GFP maintains the ability to trap the infected cell transcriptome in the nucleus, proving for the first time that mRNA degradation is not a prerequisite for vhs effects on the nuclear transcriptome. This virus has therefore uncoupled the nuclear retention and degradation activities of vhs, providing new understanding of vhs during infection.

## Introduction

The UL41 gene of herpes simplex virus 1 (HSV-1) encodes the virion host shutoff (vhs) protein, an important virulence factor that is conserved across the *alphaherpesvirinae* sub-family (1). vhs is an endoribonuclease that induces the degradation of mRNA but not rRNA or tRNA (2–4), by binding to the cellular translation initiation machinery through the eIF4A and eIF4H components of the eIF4F cap-binding complex, resulting in mRNA cleavage (1, 5–7). This activity effectively shuts down cellular protein synthesis (8, 9), freeing up ribosomes to translate viral mRNA. One consequence of vhs activity is that cellular transcripts encoding interferon-stimulated genes (ISGs), generated in response to virus infection, are targeted for degradation, thereby blunting the host innate response (10, 11). More recently, vhs has also been shown to induce the degradation of double-stranded RNA contributing to the inhibition of the protein kinase R (PKR) response seen in virus infection and revealing an additional role for vhs in counteracting host defences (12).

In theory, vhs should target all translating transcripts equally, both host and viral. However, transcriptomic analyses from our group have demonstrated that cellular mRNAs exhibit a wide range of susceptibility to vhs activity, ranging from almost complete resistance to 1000-fold reduction, suggesting that specificity for vhs activity exists (10); this result was also confirmed in another recent study (13). In the case of virus transcripts, vhs has been shown to play a key role in regulating viral gene expression during the classical herpesvirus cascade, by reducing the levels of immediate-early (IE) and early (E) viral transcripts at the transition to late (L) gene expression (2, 14). In addition, previous work from our group has revealed a novel mechanism whereby vhs causes the nuclear retention of mRNAs, providing an alternative mechanism of translation inhibition (10, 15). In particular, using mRNA FISH, IE and E transcripts were seen to be retained in the nucleus in a vhs-dependant fashion from around 10 h onwards, whereas L transcripts were efficiently exported to the cytoplasm (15). In the absence of vhs, all viral transcripts were cytoplasmic.

Concomitant with this nuclear retention of mRNA, the steady state localisation of the polyA binding proteins PABPC1 is altered from cytoplasmic to nuclear in HSV1 infected cells (10, 15, 16). PABPC1 binds the polyA tails of mRNA transcripts in the nucleus and shuttles between the nucleus and cytoplasm as mRNAs undergo nuclear export, translation and normal cellular degradation (17, 18). In uninfected cells, PABPC1 has a steady-state cytoplasmic localisation, however upon infection with HSV-1, or expression of vhs by transient transfection, PABPC1 accumulates in the nucleus in a vhs-dependant manner (10, 15), meaning that relative PABPC1 localisation is a useful surrogate for vhs function. These observations follow those seen previously for the Kaposi’s sarcoma herpesvirus (KSHV) SOX protein (19, 20), revealing the potential for a universal mechanism across the herpesviruses for regulating RNA metabolism.

Given its profound activity on the cellular transcriptome, vhs would be expected to be lethal to the virus if left unchecked, and in recent years it has become clear that the virus uses several mechanisms to lessen the consequences of vhs activity. First, the vhs transcript is inherently untranslatable, a feature which has been attributed to the presence of transferable inhibitory sequences within the mRNA (15). Second, the vhs transcript itself is predominantly localised in the nucleus when expressed during virus infection or in isolation, providing another layer of translation inhibition (15). A third process for regulating vhs activity in virus infection is through the formation of a trimeric complex between the vhs protein and the viral proteins VP16 and VP22 (15, 21), in which VP16 binds directly to vhs (22) and VP22 binds directly to VP16 (23). As such, both VP16 and VP22 deletion viruses exhibit profound shutoff of translation in the infected cell (10, 24–26), while Δ22 viruses are rescued by spontaneous mutations in vhs (10, 24, 27). This originally led to the hypothesis that vhs endoribonuclease activity is overactive in the absence of VP22, but our recent work has shown that the main outcome of deleting VP22 is not increased mRNA degradation *per se*, but increased nuclear retention of the transcriptome (10). As such, the nuclear export of L transcripts and subsequent late protein synthesis was dependent on the presence of VP22 (10).

To date, our cell biological studies of vhs have been impeded by the lack of an effective anti-vhs antibody for downstream studies. Here we report the generation and characterisation of a recombinant HSV1 which expresses vhs tagged with GFP at its C-terminus (vhs-GFP). We show that HSV1 vhs-GFP maintains many characteristics of Wt virus, including similar plaque size, viral gene expression, the ability to form a complex with VP16 and VP22 and virion packaging of vhs itself. We also show that when first expressed, vhs-GFP concentrates in a juxtanuclear position in close proximity to but not associated with the Golgi apparatus, while later in infection it co-localises with VP16. Despite vhs-GFP retaining activity in transient transfection, it failed to induce mRNA degradation or PABPC1 relocalisation to the nucleus in the context of virus infection, whereas PKR phosphorylation was enhanced, suggesting that vhs enzymatic activity is abrogated in this virus. Nonetheless, mRNA FISH of virus transcripts revealed that nuclear retention of IE and E mRNA was maintained in cells infected with HSV1 vhs-GFP, despite a lack of mRNA degradation. This unexpected result uncouples the endoribonuclease and nuclear retention activities of vhs, providing us with a unique tool to investigate these discrete molecular aspects of vhs function.

## Results

### Vhs tagged with GFP at its C-terminus retains endoribonuclease activity

To determine if vhs retains activity when fused to GFP, we first tagged vhs at either its N- or C-terminus and expressed it by transient transfection in HeLa cells. Total protein analysed by SDS-PAGE and Western blotting for GFP revealed that as expected, GFP accumulated to high levels in transfected cells, but GFP-vhs was around 15-fold lower, while expression of vhs-GFP was not detectable (Fig 1A). These results are in line with previous experiments by us and others where it was shown that the vhs transcript is inherently untranslatable, and vhs protein fails to accumulate during transient transfection despite vhs mRNA levels being similar to those in infected cells (1, 15, 21). The ability of GFP-vhs or vhs-GFP to relocalise PABPC1 to the nucleus of HeLa cells, indicative of endoribonuclease activity, was measured using immunofluorescence of PABPC1. Following transfection with a plasmid expressing GFP alone, PABPC1 remained cytoplasmic (Fig 1B). However, while vhs-GFP caused PABPC1 relocalisation to the nucleus, GFP-vhs failed to do so, indicating that vhs functionality is retained only in vhs tagged at its C-terminus with GFP (Fig 1B). Furthermore, mRNA FISH using a probe for GFP indicated that while the GFP and GFP-vhs transcripts were localised in the cytoplasm, the vhs-GFP transcript was retained in the nucleus (Fig 2). This confirms that unlike GFP-vhs, vhs-GFP seems to retain the full characteristics of the native vhs transcript (15).

**FIG 1.**
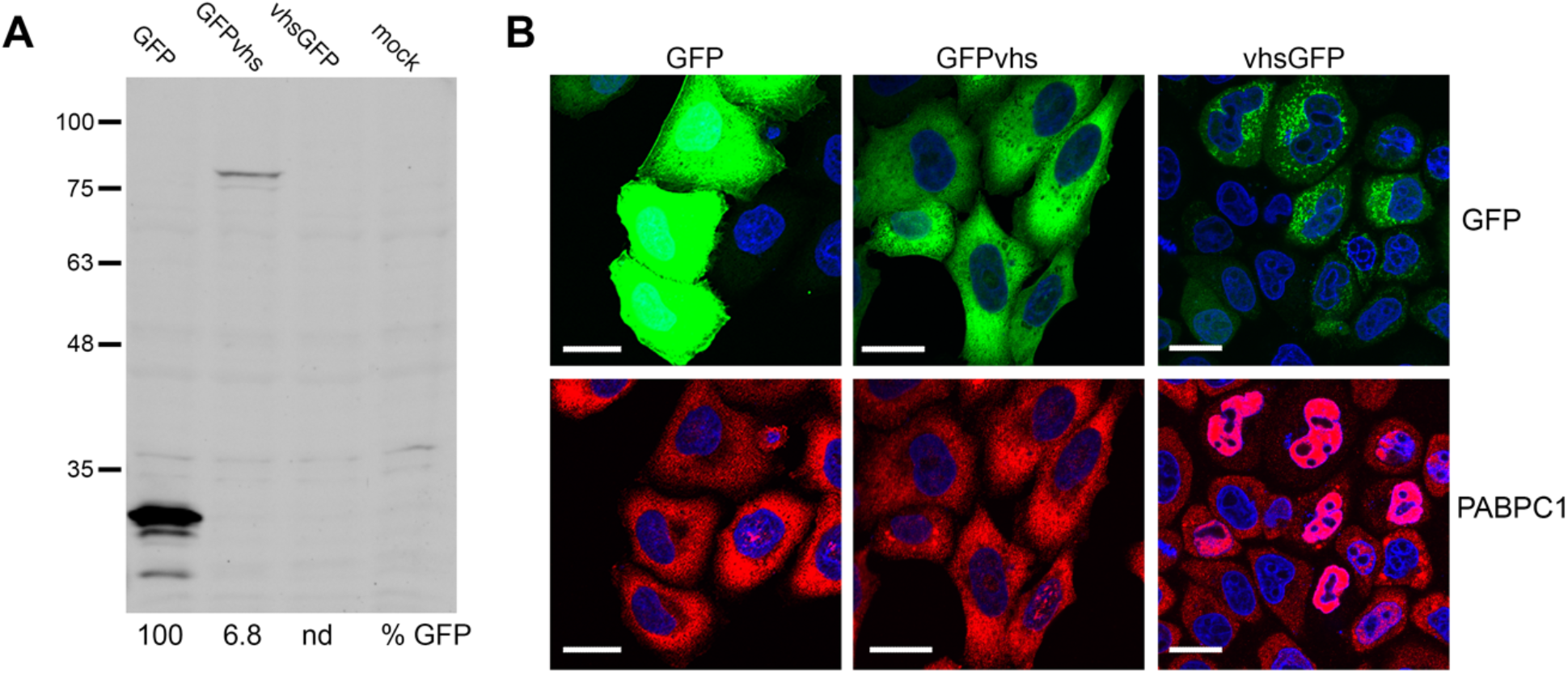
Expression of GFP-tagged vhs proteins. **(A)** HeLa cells were mock transfected or transfected with plasmids expressing GFP-tagged proteins as shown, and after 20 h total cell lysates were analysed by Western blotting for GFP. Relative expression of GFP was quantitated using LICOR ImageStudio and is represented as percentage of untagged GFP level. **(B)** As for (A), but transfected cells grown on coverslips were fixed and stained for PABPC1 (red) and nuclei stained with DAPI (blue). Scale bar = 20 μm.

**FIG 2.**
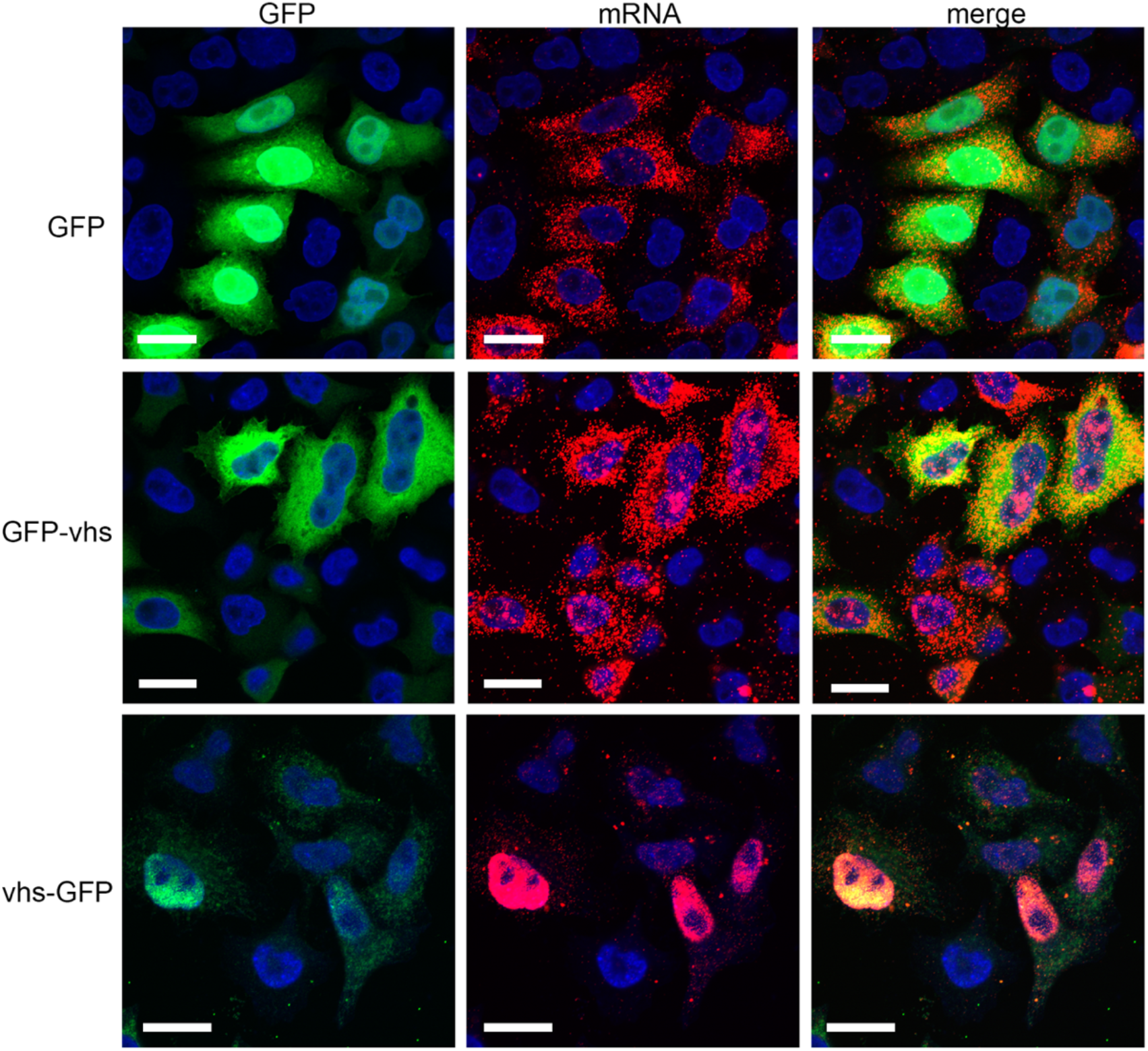
Nuclear localisation of the vhs-GFP transcript in transfected cells. HeLa cells grown in two-well chamber slides were transfected with plasmids expressing GFP-tagged proteins as shown, and after 20 h were fixed in 4% paraformaldehyde. Cells were then processed for mRNA FISH using a GFP-specific probe (red) and nuclei stained with DAPI (blue). GFP fluorescence is in green. Scale bar = 20 μm.

### Characterisation of recombinant HSV1 expressing a vhs-GFP fusion protein

A recombinant HSV1 in strain Sc16 was subsequently generated using homologous recombination to express vhs-GFP in place of vhs (Fig 3A). Following three rounds of plaque purification on Vero cells, vhs-GFP expression was confirmed by analysing total infected cell protein by SDS-PAGE and Western blotting for GFP, vhs, and a loading control of α-tubulin (Fig 3B). Another recombinant virus expressing a different tagged protein – YFP-UL47 - was analysed simultaneously to directly compare the relative expression of vhs with a second tagged virus protein (Fig 3B). In HSV1-vhs-GFP infected cells, a protein band of the correct size was detected with both the vhs and the GFP antibodies, confirming the expression of vhs-GFP in this virus (Fig 3B). Moreover, while the vhs antibody indicated that vhs was expressed at roughly the same level in cells infected with HSV1 expressing either the vhs-GFP or YFP-UL47, quantification of the respective vhs-GFP and YFP-UL47 bands using the GFP antibody (which detects YFP with equal efficiency), indicated that vhs is expressed at a level 15-fold lower than UL47, re-enforcing the relatively poor expression of vhs (Fig 3B). Plaque assays on Vero and HFFF cells further indicated that the plaque size of HSV1-vhs-GFP was similar to that of the Wt Sc16 virus suggesting that virus replication was not significantly impaired by the expression of vhs-GFP in place of vhs (Fig 3C). The growth characteristics of vhs-GFP were next compared to that of Wt strain Sc16 in a single-step growth curve in HFFF cells. Cells were infected at MOI 2 and total virus was harvested every 5 hours up to 30 h, and titrated on Vero cells, indicating that vhs-GFP replication was delayed by around 5 hours compared to Wt virus but ultimately reached approximately the same titre (Fig 3D). A similar infection time-course was analysed for protein expression by SDS-PAGE and Western blotting to assess the kinetics of expression of representative viral proteins (Fig 3E). Blotting for ICP27, TK and VP16, representative of the viral gene classes IE, E, and L respectively, showed that the kinetics of expression of all three classes was similar between Wt and HSV1-vhs-GFP infections, while vhs and GFP blots confirmed the expression of vhs-GFP with similar kinetics to vhs alone, although vhs-GFP accumulated to lower levels than vhs (Fig 3E).

**FIG 3.**
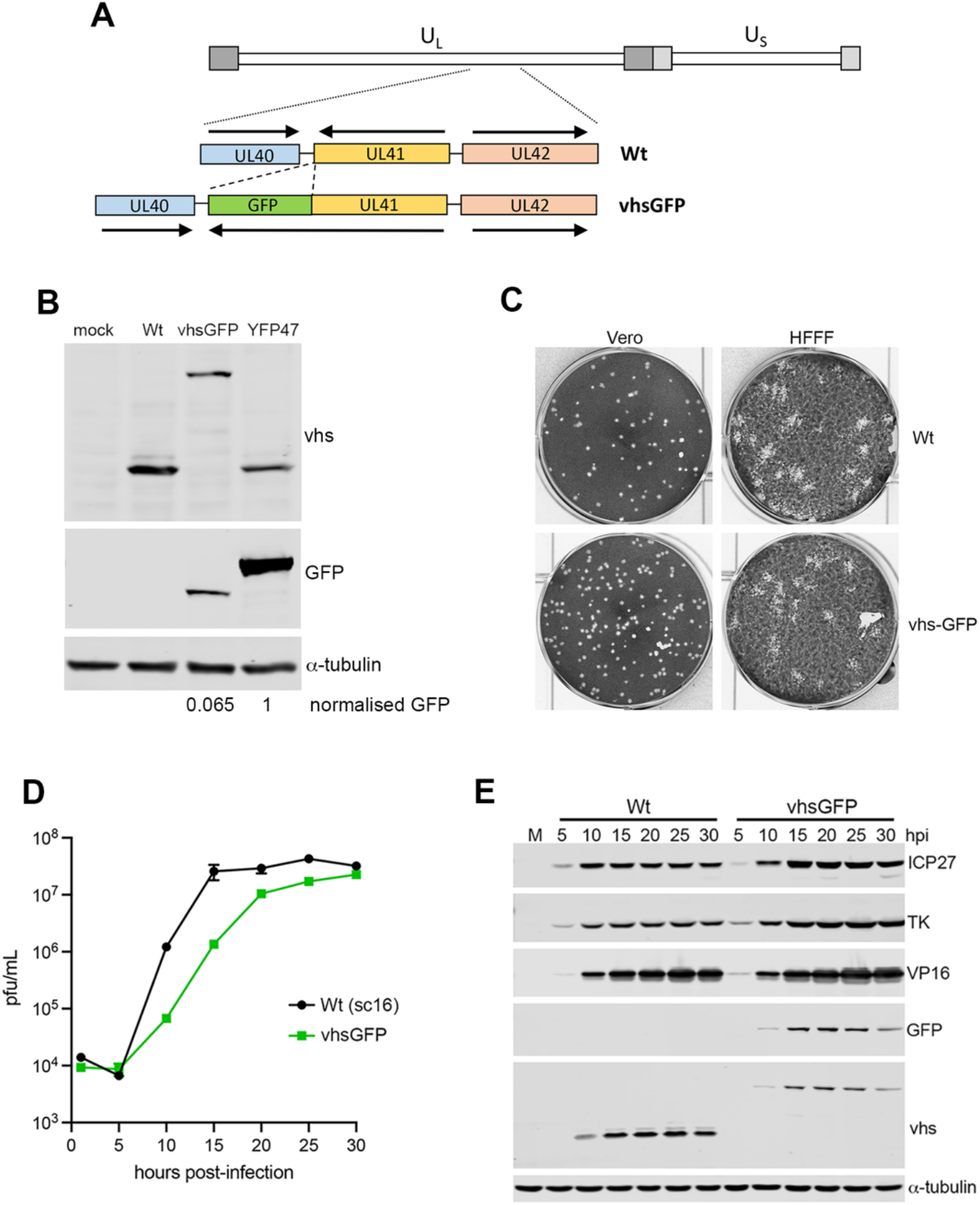
Construction and characterisation of HSV1 expressing vhs-GFP. **(A)** HSV1 expressing vhs-GFP in place of GFP was constructed by co-transfecting Vero cells with infectious Sc16 genomic DNA and plasmid containing the UL41GFP fusion gene surrounded by the flanking sequences from UL40 and UL42. **(B)** HFFF cells were mock infected or infected with Sc16, plaque purified vhs-GFP or YFP-UL47 viruses at MOI 2, and total protein harvested at 16 hpi prior to separation by SDS-PAGE and Western blotting with antibodies for GFP, vhs and α-tubulin. **(C)** Vero and HFFF cells were infected with around 40 pfu of Sc16 or vhs-GFP and incubated for three days prior to fixation and staining with crystal violet. **(D)** One-step growth curves for Wt (Sc16) and HSV1 vhs-GFP viruses were carried out by harvesting total virus from HFFF cells infected at MOI 2 every 5 hours up to 30 hours. Samples were titrated onto Vero cells. The mean titre of three replicates and associated SEM are plotted. **(E)** Total extracts of HFFF cells infected as in (D) were analysed by SDS-PAGE and Western blotting for the indicated virus proteins and α-tubulin as a loading control.

To investigate where vhs localises in the infected cell, HFFF cells infected with HSV1 vhs-GFP were fixed at various times after infection and imaged for GFP fluorescence using confocal microscopy. In line with Western blotting, vhs-GFP was first detected at low levels at around 10 hours and increased in intensity to 15 hours, when it was particularly concentrated close to the nucleus and at the cell periphery (Fig 4A). Closer examination of the 10-hour sample revealed that vhs-GFP was clustered in a juxtanuclear location when first expressed at detectable levels (Fig 4B), but staining of cells at this time with an antibody for giantin indicated that these clusters, while in proximity of the Golgi apparatus, were not associated with it (Fig 4C).

**FIG 4.**
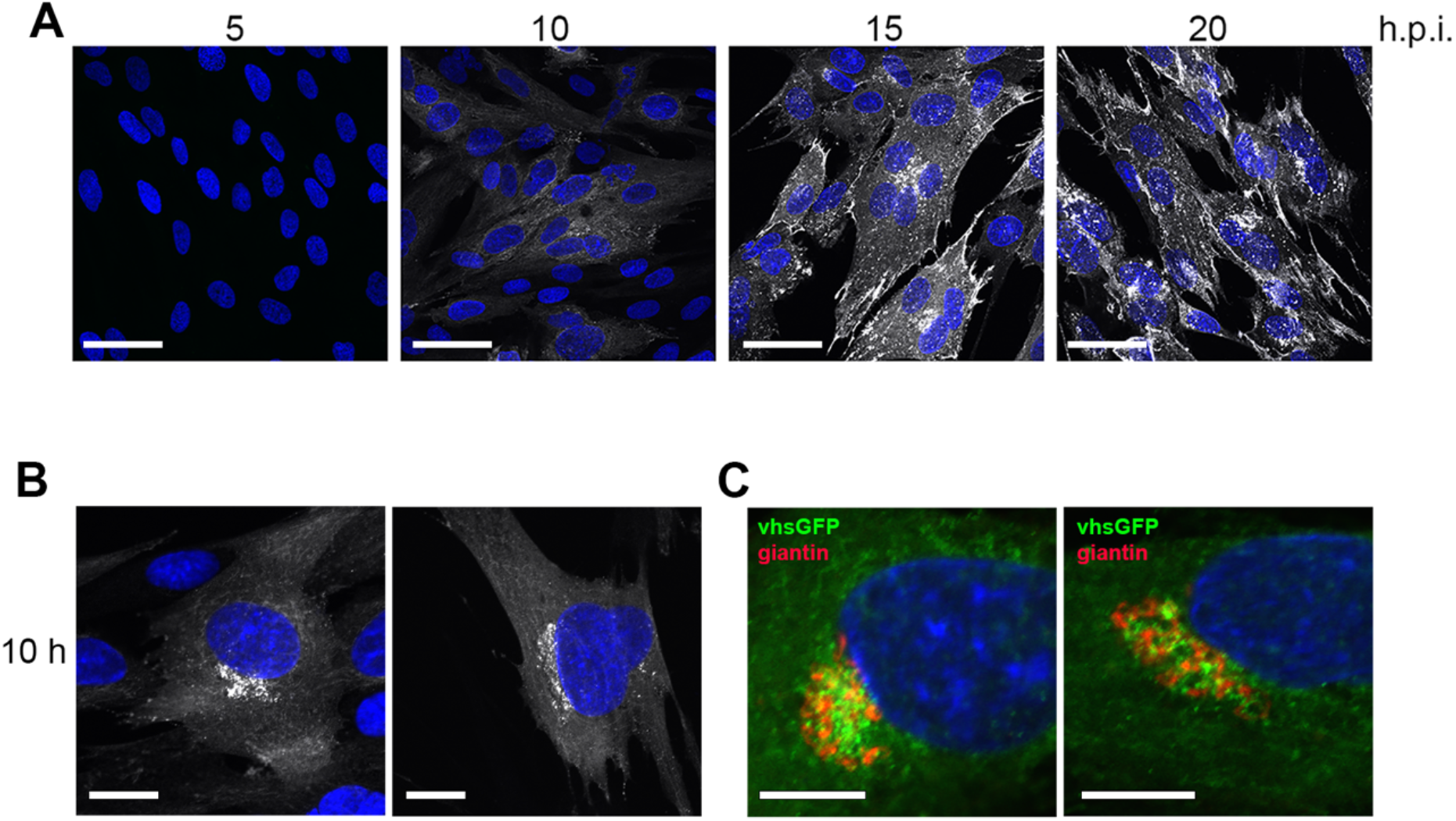
Localisation of vhs-GFP during HSV1 infection. **(A)** HFFF cells infected with HSV1-vhs-GFP MOI 2 were fixed at the indicated times, stained with DAPI (blue) and imaged for GFP fluorescence (white) using confocal microscopy. Scale bar = 50 μm. **(B)** Examples of the 10 h sample from (A) imaged at higher magnification. Scale bar = 20 μm**. (C)** HFFF cells infected as in (A) were fixed and stained for the Golgi marker giantin. Scale bar = 10 μm.

### The vhs-GFP fusion protein maintains the ability to form the vhs-VP16-VP22 complex in infected cells

Viral protein VP16 binds directly to vhs, and together with VP22 regulates the activity of vhs (22, 26). Immunofluorescence of VP16 in HSV1 vhs-GFP infected cells indicated that these two proteins were extensively co-localised at later times in infection when virus assembly would be optimal (Fig 5A). To determine if vhs-GFP interacts with VP16 and VP22 during infection, a GFP-TRAP pulldown was performed on extracts of HaCat cells infected with either vhs-GFP or Wt viruses at MOI 5, harvested at 24 h. The resulting pulldown complex was analysed by Western blotting, indicating that both VP16 and VP22 were present in the vhs-GFP interactome but not the negative control (Fig 5B), suggesting that the fusion protein is capable of binding its usual viral protein partners.

**FIG 5.**
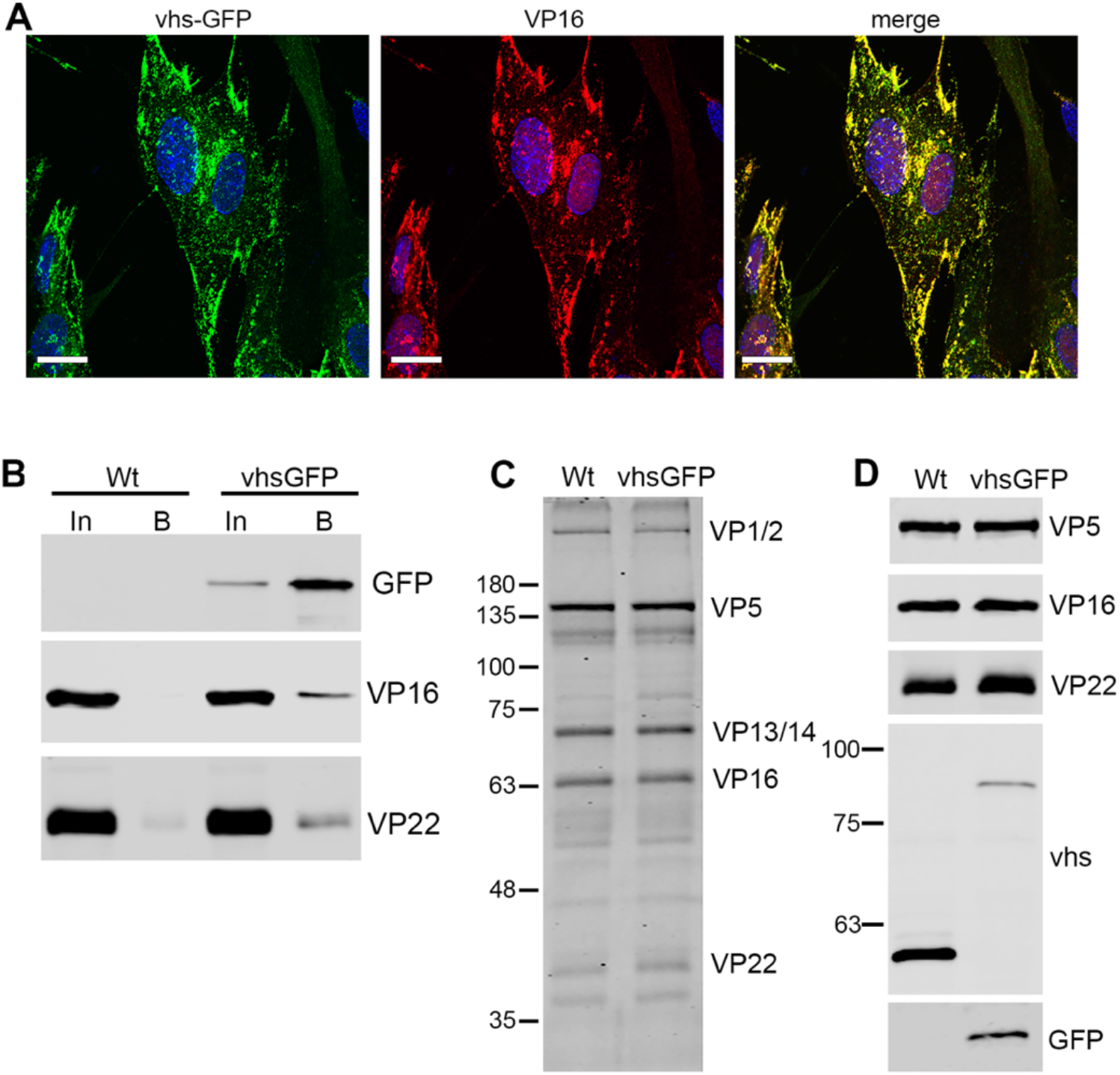
Interaction of vhs-GFP with VP16 and assembly into virions. **(A)** HFFF cells infected with HSV1 vhs-GFP MOI 2 were fixed at 16 h, stained for VP16 (red) and imaged for GFP (green). Nuclei were stained with DAPI (blue). Scale bar = 20 μm. **(B**) GFP-TRAP pulldown was carried out on HaCat cells infected with Wt (Sc16) or vhs-GFP at moi 5 and harvested 24 hpi. The pulldowns were analysed by SDS-PAGE and Western blotting with antibodies to GFP, VP16 and VP22. In = input, B = pulldown **(C) & (D)** Extracellular virions were purified on 5 – 15% Ficoll gradients and analysed by SDS-PAGE followed by Coomassie blue staining (C) or Western blotting with antibodies as indicated (D).

The interaction between VP16 and vhs has also been implicated in the packaging of vhs into the virion (22). To test the assembly of vhs-GFP, extracellular HSV1 vhs-GFP and Wt virions were purified from infected HaCat cells by centrifuging on a 5 to 15% Ficoll gradient. Solubilised virions were analysed by SDS-PAGE followed by Coomassie blue staining and Western blotting (Fig 5C and 5D). Using the VP5 major capsid protein band to equalise virion loading, the total protein profiles for each virus was determined to be comparable (Fig 5C). Western blotting for viral proteins vhs, GFP, VP16 and VP22, using VP5 as a loading control confirmed the presence of the ~55 kDa vhs protein in Wt virions and the ~82 kDa fusion protein in vhs-GFP virions (Fig 5D). Although VP16 was equally packaged in both virion samples, vhs-GFP was present around five-fold lower than Wt vhs (Fig 5D) suggesting that packaging of vhs-GFP into virions may be less efficient than Wt virus.

### vhs-GFP lacks endoribonuclease activity in infected cells

We previously showed that PABPC1 was relocalised from the cytoplasm to the nucleus in a vhs-dependent manner in both HSV1 infected cells and in cells transfected with a plasmid expressing vhs (22). During the initial transfection experiments we also showed that when transfected into Hela cells, vhs-GFP was also capable of relocalising PABPC1 to the nucleus (Fig 1B). To confirm that this activity was maintained in HSV1 vhs-GFP, HFFF cells infected with either Wt or HSV1 vhs-GFP were fixed at 16 h and stained for PABPC1. As shown previously, PABPC1 was concentrated in nuclear accumulations in Wt infected cells (Fig 6). By contrast, PABPC1 remained cytoplasmic in HSV1 vhs-GFP infected cells even as late as 20 h after infection (Fig 6).

**FIG 6.**
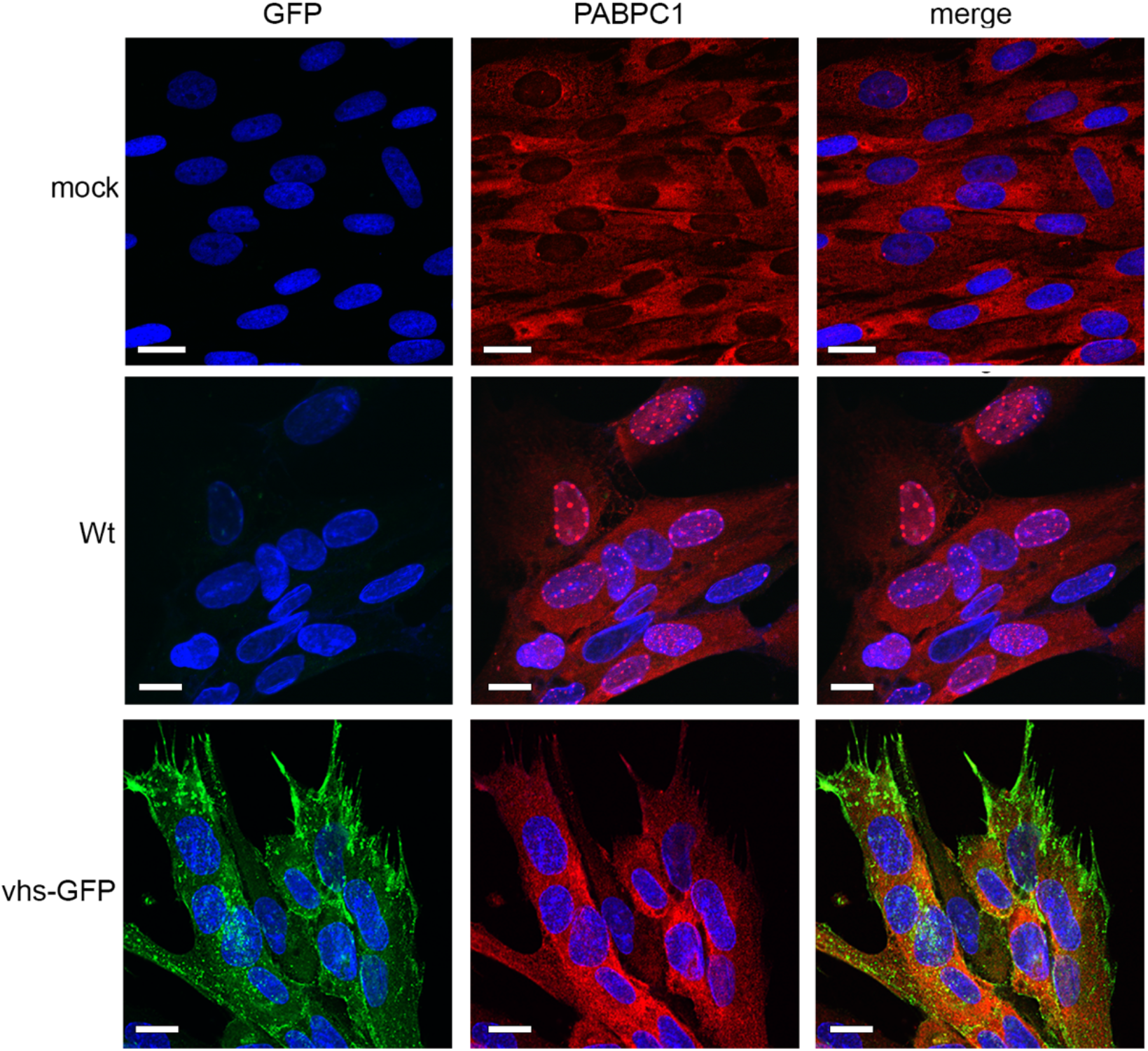
PABPC1 is not relocalised in HSV1 vhs-GFP infection. HFFF cells infected with Wt (Sc16) or HSV1 vhs-GFP viruses at MOI 2 were fixed at 16 hours, stained with an antibody for PABPC1 (red) and nuclei stained with DAPI (blue). Scale bar = 20 μm.

The lack of PABPC1 relocalisation to the nucleus raised the possibility that vhs may not be active as an endoribonuclease when expressed in the context of virus infection. To test the ability of the vhs-GFP fusion protein to degrade cellular mRNA, relative levels of two ISG transcripts (IFIT1 and IFIT2) and two transcripts previously shown to be hypersensitive to vhs activity (MMP1 and MMP3) were measured by RT-qPCR at 16 h. IFIT1 and IFIT2 transcripts were reduced in Wt infection compared to mock-infected control (Fig 7A) as shown in our previous work (10). By contrast these ISG transcripts were both upregulated in HSV1 vhs-GFP infection by around Log_2_ fold change of 3, suggesting that these transcripts were not degraded by vhs-GFP and were being activated in response to virus infection, as shown previously in cells infected. With a vhs knockout virus (10). In the case of MMP1 and MMP3, these transcripts were reduced by Log_2_ fold change of 8 in Wt infection, again as shown previously (10), but these transcripts were unchanged in the HSV1 vhs-GFP infection compared to the mock-infected control at 16 h (Fig 7A). For all four transcripts, very similar results were observed at 24 h to those seen at 16 h, suggesting that the ability of vhs-GFP to degrade cellular transcripts is impaired, rather than delayed.

**FIG 7.**
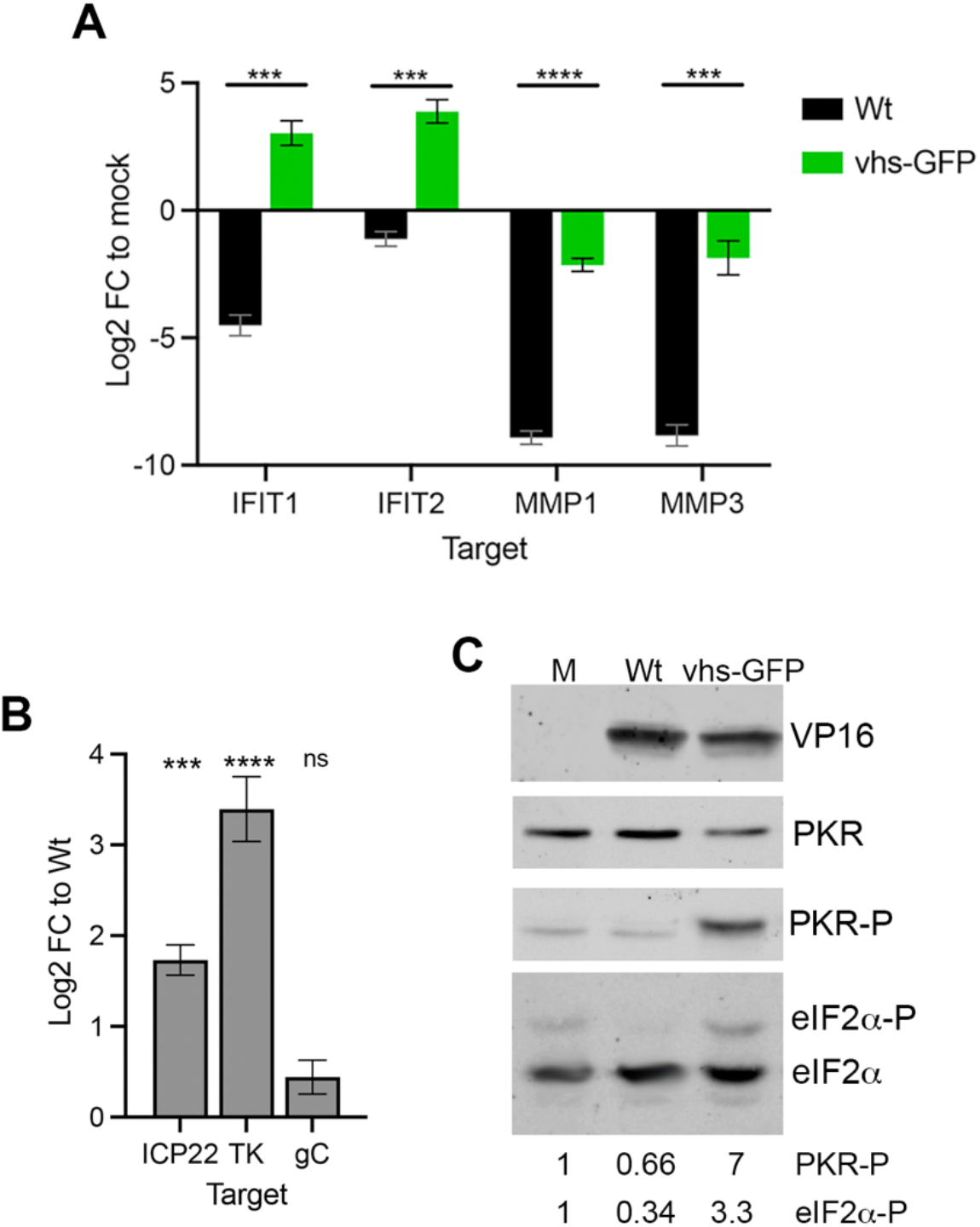
vhs-GFP expressed in virus infection is deficient in endoribonuclease activity. **(A)** HFFF cells were infected with Wt or HSV1 vhs-GFP viruses at MOI of 5. Total RNA was purified at 16 hpi and subjected to RT-qPCR for cellular transcripts previously identified as being susceptible to vhs activity. **(B)** RNA samples from (A) were subjected to RT-qPCR of viral transcripts ICP22 (IE), TK (E) and gC (L). The mean and ± standard error for n = 3 is shown. Statistical analysis was carried out using an unpaired t test. ***, p < 0.001. ****, p < 0.0001. ns, not significant. **(C)** HFFF cells infected with Wt or HSV1 vhs-GFP viruses at MOI of 2 were harvested at 16 h and analysed by SDS-PAGE and Western blotting with antibodies as indicated. Western blotting for eIF2α was carried out using 10μm Phos-Tag PAGE. Relative level of phospho-PKR and phospho-eIF2α was quantitated using LICOR Image Studio and is represented relative to the level in uninfected cells.

Viral transcripts representing IE (ICP22), E (TK) and L (gC) genes were measured in the same way as the cellular transcripts, but in this case the log_2_ FC was compared to Wt infected cells. ICP22 and TK transcripts were significantly more abundant in HSV1 vhs-GFP infection compared to Wt (Fig 7B), but the level of the late gC transcript was not significantly different between the two viruses, in agreement with our previous results on Δvhs infected cells (10). Hence, the significant differences in IE and E mRNA transcript levels between HSV1 vhs-GFP and Wt infection further suggests that the vhs-GFP fusion protein is severely impaired or non-functional with respect to its mRNA degradation capabilities.

One other consequence of vhs endoribonuclease activity is the degradation of double-stranded RNA, formed by the annealing of virus mRNA transcribed from both strands of the virus genome, thereby contributing to the suppression of PKR phosphorylation and subsequent eIF2α phosphorylation in infected cells to subsequently counteract translational shutoff (12). This suggests that vhs acts in tandem with two other virus factors: Us11, which blocks PKR phosphorylation (28) and ICP34.5, which controls eIF2α phosphorylation (29). Western blotting of Wt and HSV1 vhs-GFP infected cell extracts indicated that PKR phosphorylation was indeed enhanced in cells expressing vhs-GFP compared to Wt virus infection (Fig 7C). Increased PKR phosphorylation was also reflected in an increase in eIF2α phosphorylation in HSV1 vhs-GFP infection.

### Nuclear retention of the infected cell transcriptome in HSV1 vhs-GFP infected cells

The failure of vhs-GFP to degrade infected cell mRNA led us to hypothesize that this virus would be equivalent to a Δvhs virus in that all vhs activities would be abrogated by the fusion of GFP at its C-terminus in the context of virus infection. As we have not yet formally shown that vhs endoribonuclease activity is required for the nuclear retention of the infected cell transcriptome (10, 15), mRNA FISH was carried out on infected HFFF cells for representative IE (ICP27) E (TK) and L (gD) transcripts. Wt infected cells recapitulated our previous results that IE and E transcripts but not L transcripts were retained in the nucleus at later times of infection (Fig 8A, Wt), but in Δvhs infected cells, all classes of transcript were entirely cytoplasmic, thereby confirming that vhs is required for nuclear retention of these transcripts (Fig 8A, Δvhs). However, in the HSV1 vhs-GFP infected cells, IE and E transcripts were also seen to be retained in the nucleus (Fig 8A, vhs-GFP), despite vhs-GFP failing to induce the degradation of mRNA or the nuclear retention of PABPC1. Likewise, the cellular transcript for MMP1, which is efficiently degraded by vhs but not vhs-GFP (Fig 7A), was also retained in the nucleus despite not being degraded in HSV1 vhs-GFP infected cells, in contrast to Δvhs infected cells (Fig 8B). Finally, the localisation of the vhs-GFP transcript itself was examined in infected cells using a probe for GFP or vhs. This revealed that as for Wt infection, where the vhs transcript was predominantly nuclear, the vhs-GFP transcript was also retained in the nucleus (Fig 8C), confirming the results obtained above in cells expressing vhs-GFP by transfection.

**FIG 8.**
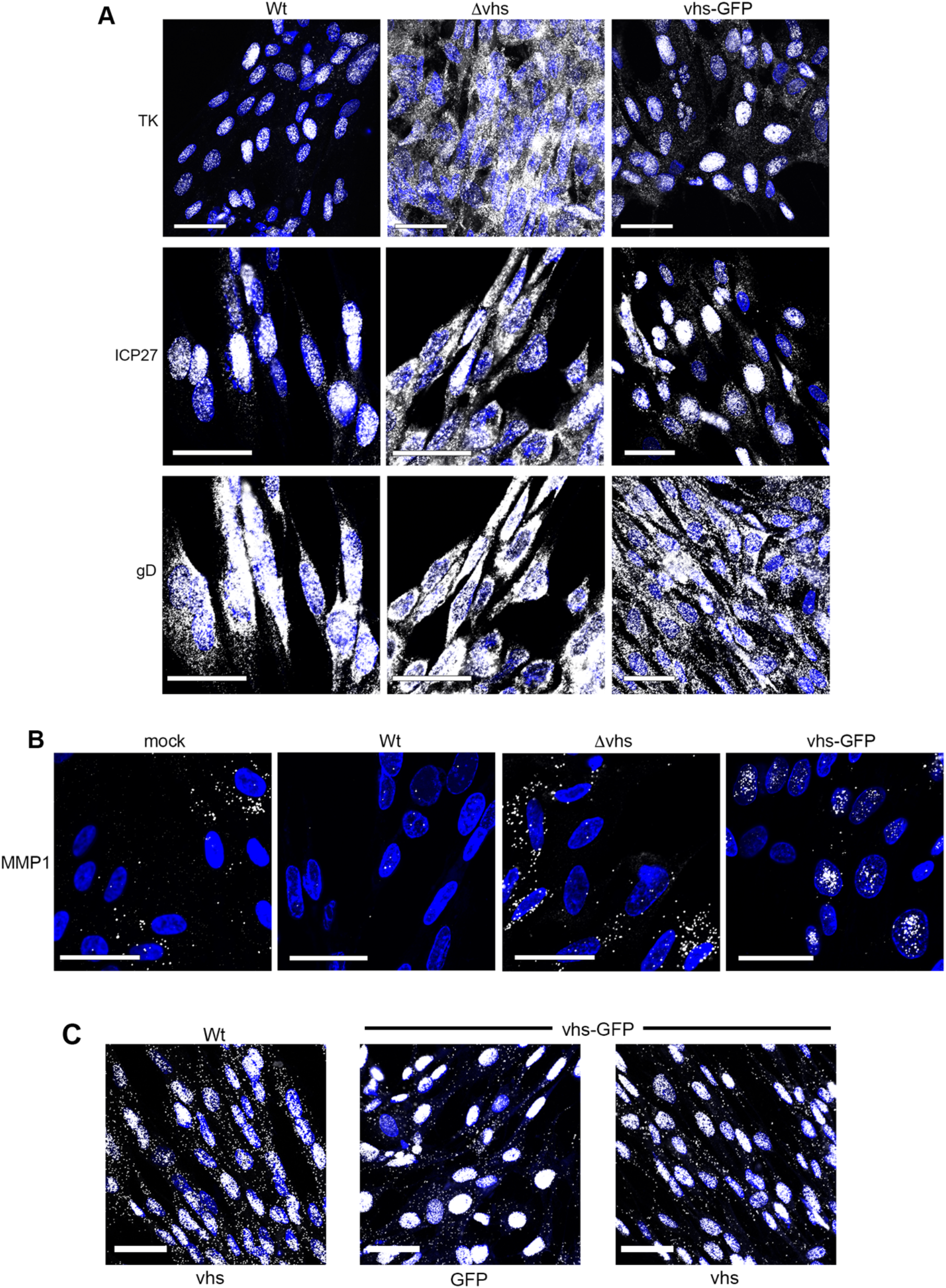
vhs-GFP maintains the ability to cause the nuclear retention of viral and cellular transcripts. HFFF cells grown in two-well slide chambers were infected with Wt, Δvhs or HSV1 vhs-GFP MOI 2 and fixed after 16 hours in 4% paraformaldehyde. Cells were then processed for mRNA FISH using probes specific for (**A)** IE (ICP27), E (TK) and L (gD) transcripts or (**B)** MMP1 (all in white). NB The same field in Wt and Δvhs infection is shown for ICP27 and gD mRNA FISH. (**C)** As above but Wt or HSV1 vhs-GFP infected cells were stained with a probe for UL41 or GFP (in white). Nuclei were stained with DAPI (blue). Scale bar = 50 μm.

These unexpected results indicate that although vhs-GFP lacks endoribonuclease activity in HSV1 infected cells, it retains the ability to alter the compartmentalisation of the infected cell transcriptome, both viral and cellular. Hence, these two properties of vhs have been uncoupled in the HSV1 vhs-GFP virus, providing scope for further molecular understanding of vhs activity.

## Discussion

The vhs protein of HSV1 is involved in regulating the RNA environment of the infected cell. Not only does it induce the degradation of mRNA and dsRNA (3, 4, 12), but it also causes the nuclear retention of both the viral and the cellular transcriptome (10). The overall effect of this activity is to enhance translation of the late virus transcriptome not only by sequestration of the early virus transcriptome in the nucleus, but also by reducing the PKR stress response (12, 30). As a virulence factor, vhs therefore helps to optimise the virus infection process and block the establishment of an antiviral environment in the cell. In this study we aimed to develop a cell biology tool for studying vhs, using a vhs-GFP fusion protein which had been demonstrated to function when expressed in isolation. Having constructed recombinant HSV1 expressing vhs-GFP in place of native vhs, we characterised its localisation, binding partners, virion packaging, and endoribonuclease activity during infection. Surprisingly, despite functioning in isolation and maintaining its physical characteristics during infection, vhs-GFP exhibited little if any endoribonuclease activity, a result that was backed up by the failure of PABPC1 to relocalise to the nucleus. This suggests that the addition of GFP to the C-terminus of vhs differentially affects its activity depending on whether it is expressed in isolation or in virus infection.

The vhs-GFP protein was expressed at a relatively low level during transfection or infection, with a 20-fold or over reduction in GFP expression by fusing it to the vhs open reading frame at either the N- or C- terminus, (Fig 1A). This is not unexpected as we have previously demonstrated that vhs autoregulates its own expression by various means, including the nuclear retention of its own transcript, and inefficient translation of its transcript (15). Despite the low level of vhs expression during infection, the HSV1 vhs-GFP virus nonetheless afforded us with the opportunity to investigate vhs localisation throughout the virus infection cycle. The vhs-GFP fusion protein was only detectable at 8 to 10 hours after infection, at which time it localised to juxtanuclear clusters that were close to but not associated with the Golgi apparatus. As infection progressed, fluorescence levels increased and vhs-GFP appeared throughout the cytoplasm and at the cell periphery, where it colocalised with its known binding partner, VP16. As both vhs and VP16 are assembled in to the HSV1 virion, these sites of vhs positivity are likely to represent different stages of the HSV1 morphogenesis pathway including the trafficking of structural proteins on the path to virus envelopment and the egress of enveloped virions (31).

Herpesvirus encoded endoribonucleases act by enhancing the normal rate of turnover of cellular mRNA; mRNA is first cleaved by the endoribonuclease before the cellular mRNA decay machinery acts on it (32, 33). This accelerated cleavage alters the steady-state localisation of PABPC1 from cytoplasmic to nuclear as it shuttles between the nucleus and the cytoplasm (17). The most striking result from this study is that despite the lack of endoribonuclease activity and PABPC1 retention in the nucleus, vhs-GFP maintains the ability to cause the vhs-dependent nuclear retention of IE and E transcripts seen in Wt infection. This reveals for the first time that vhs cleavage and degradation of mRNA is not a prerequisite for enhanced nuclear compartmentalisation of the infected cell transcriptome. It also emphasizes that the relative steady-state localisation of PABPC1 is a direct consequence of vhs endoribonuclease activity and mRNA degradation in the cytoplasm, rather than a correlate for the relative compartmentalisation of the transcriptome: PABPC1 bound to the polyA tails of cytoplasmic transcripts can only return to the nucleus after mRNA degradation has released it for nuclear import (34). Much of what is known about PABPC1 compartmentalisation has been elucidated from work on virus endoribonucleases, and in particular the SOX protein from Kaposi’s sarcoma herpesvirus (KSHV). As for vhs, the activity of SOX in the cytoplasm causes the nuclear retention of both PABPC1 and mRNA (19), but in the case of SOX, previous work has revealed that the inhibition of mRNA export is a direct consequence of the nuclear accumulation of PABPC1 (35). However, our results shown here demonstrate that vhs induces the nuclear accumulation of mRNA without concomitant relocalisation of PABPC1, showing not only that these two properties of vhs are separable, but also that PABPC1 accumulation in the nucleus is not a direct consequence of increased nuclear concentration of mRNA (36).

The molecular explanation for the vhs-induced sequestration of the transcriptome in the nucleus will require further studies, but one potential explanation is that vhs causes the cytoplasmic retention of a nuclear export factor, preventing it from returning to the nucleus and thereby blocking the subsequent export of IE and E transcripts. If this is the case then it raises the question of how L transcripts are successfully exported for translation of structural proteins. The activities of several other virus proteins may provide clues to this. First, the IE protein ICP27 is known to be essential for late protein expression (37, 38). ICP27 is a nucleocytoplasmic shuttling protein that binds mRNA transcripts in the nucleus and helps export them in co-ordination with the TREX (transcription-coupled export) complex, specifically through binding to the Aly/REF factor within the complex (39, 40). ICP27 may therefore counteract the nuclear retention activity of vhs by bypassing the need for any specific factors retained by vhs in the cytoplasm. Interestingly, ICP27 has been shown to stimulate the translation of a subset of late HSV1 mRNAs (41), a property likely to be explained by its ability to directly bind PABPC1 and recruit it to these mRNAs (42). Additionally, deletion of either of the VP16 and VP22 components of the vhs-VP16-VP22 complex causes translational shut-off, and in the case of VP22 at least, this shutoff correlates with the entrapment of late transcripts in the nucleus (10). Hence, VP22 and VP16 quench the ability of vhs to retain the nuclear transcriptome, potentially by competing for vhs binding to specific factors in the cytoplasm, to function in co-ordination with ICP27 to activate late protein expression. GFP-Trap analysis of vhs-GFP expressed during infection will now provide us with the means of investigating the binding partners of vhs during infection, allowing us to investigate the full vhs interactome and identify potential candidates required for export of viral mRNAs late in infection.

These results further emphasize the role that vhs plays in regulating the antiviral response to HSV1 infection. As we have shown before, ISG transcripts are initially upregulated in HSV1 infected HFFF, before dropping again as infection progresses, with this decrease in ISG level being dependent on vhs activity (10). Here we have shown that vhs-GFP also fails to degrade IFIT1 and IFIT2 transcripts, confirming the dependence of this process on vhs endoribonuclease activity. The PKR stress response is also upregulated in HSV1 vhs-GFP infection with a seven-fold increase in PKR phosphorylation and a 3.5-fold increase in eIF2α phosphorylation compared to Wt infection, suggesting that the level of dsRNA may be increased in these infected cells. Nonetheless, there was no evidence of translational shutoff in the HSV1 vhs-GFP infected cells, indicating that the neurovirulence factor ICP34.5, which induces the dephosphorylation of eIF2α (29), is likely to counteract the increase in eIF2α phosphorylation induced by the increased PKR phosphorylation seen in cells infected with virus expressing vhs-GFP. This lack of stress-induced translational shutoff is further emphasized by the fact that, despite remaining cytoplasmic in HSV1 vhs-GFP infected cells, PABPC1 did not localise to cytoplasmic foci, suggesting that recruitment of mRNA into stress granules was not a feature of this infection (43).

In combination, these results serve to further emphasize the interplay between a range of virus factors, including the dsRNA binding protein Us11 which inhibits PKR, the neurovirulence factor ICP34.5, which reverses the phosphorylation of eiF2α, and vhs, which degrades ISG transcripts and dsRNA in the antagonism of host responses to HSV infection (44). Our new virus tool that separates out vhs-induced mRNA degradation from its effects on the localisation of the virus transcriptome affords us a unique opportunity to investigate the relative contributions of these vhs activities to the success of an HSV1 infection.

## Materials and Methods

### Cells and viruses

Vero, HFFF, HaCat and HeLa cells were cultured in Dulbecco modified Eagle medium (DMEM) supplemented with 10% foetal bovine serum (FBS). Viruses were routinely propagated and titrated in Vero cells using DMEM with 2% FBS and supplemented with 5% human serum for titrations. The parental virus strains used in this study were HSV-1 strain Sc16 and s17. The s17-derived vhs knockout virus (Δvhs) has been described before (28). Extracellular virions were gradient purified as described previously (16). Briefly, ten 175-cm^2^ flasks of confluent HaCaT cells were infected at a multiplicity of 0.05. Once cytopathic effect was advanced (3-4 days post-infection), the extracellular medium was collected and centrifuged at 3,000 rpm for 30 min at 4°C in a fixed-angle rotor to remove cell debris. Virus particles were pelleted from the supernatant at 9,000 rpm for 90 min at 4°C. The particle pellet was resuspended in 0.5 mL phosphate-buffered saline (PBS) and layered onto a preformed 11 mL 5% to 15% (wt/vol) Ficoll gradient in a 13.2 mL thin-wall polyallomer ultracentrifuge tube (Beckman Coulter). Gradients were centrifuged at 12,000 rpm for 2 h at 4°C in an SW41 Ti swinging-bucket rotor in a Sorvall Discovery SE ultracentrifuge. Virions were harvested by needle puncture through the side of the tube with a 23-gauge hypodermic needle in a volume of <1 mL, diluted in 10 mL PBS, and pelleted at 25,000 rpm for 1 h at 4°C using the same rotor and ultracentrifuge. The pellets were resuspended in 100 μL PBS and stored at −80°C. Virions were solubilised in SDS-PAGE lysis buffer prior to analysis by SDS-PAGE and Western blotting.

### Plasmids

To construct pGFP-vhs, the vhs open reading frame (UL41) was transferred from plasmid V5-vhs to pEGFPC1. To construct pvhs-GFP, UL41 was amplified by PCR from plasmid pGE204 (15) using primers F-gg**<u>agatct</u>**acatgggtttgttcgggatga and R-gc**<u>gaattc</u>**tcgtcccagaattt, and inserted into pEGFPN1. Plasmids were transfected into HeLa cells with Lipofectamine 2000.

A transfer vector to produce recombinant HSV1 vhs-GFP was constructed by amplifying the 800 bp downstream flanking region of the vhs-encoding gene UL41 from the Sc16 genome as a Not1 fragment using primers F-5’cg**<u>gcggccgc</u>**cgtcagacgagcgcgcttg3’ and R-5’cg**<u>gcggccgc</u>**cgtggccggtaccatcaac3’, and inserting the amplified fragment in the Not1 site downstream of GFP in pvhs-GFP.

### Construction of vhs-GFP virus

Equal amounts of plasmid vhs-GFP and infectious HSV1 strain Sc16 DNA were transfected into 2×10^5^ Vero cells grown in a well of a 12-well culture plate using lipofectamine 2000. Five days later when cytopathic effect was extensive, the infected cells were scraped into the cell medium, subjected three times to freeze-thawing and titrated on Vero cells. Fluorescent plaques were picked and plaque-purified a further two times before further analysis.

### Antibodies

The VP22 (AGV031) antibody has been described elsewhere (29). Other antibodies used in this study were kindly provided by the following individuals: VP16 (LP1), Colin Crump (University of Cambridge); vhs, Duncan Wilson (Albert Einstein College of Medicine); TK, Frazer Rixon (Centre for Virus Research, Glasgow). Other antibodies were purchased commercially: ICP27 (AbCam); α-tubulin (Sigma), ICP5 (Virusys), PABPC1 (Santa Cruz), GFP (Clontech), PKR, eIF2α (Cell Signalling Technology), Phospho-PKR and giantin (both AbCam). Fluorescent IRDye secondary antibodies were from LI-COR.

### SDS-PAGE and Western blotting

Total protein samples were analysed by SDS-polyacrylamide gel electrophoresis (PAGE) and were either stained with Coomassie blue or transferred to a nitrocellulose membrane for Western blot analysis. Western blots were visualised on the Odyssey CLx system (LI-COR). Phosphorylated eIF2α was distinguished from eIF2α using Phos-tag SDS-PAGE (Alpha Laboratories).

### GFP-TRAP pulldown assay

A GFP-TRAP agarose kit (Chromotek) was used to purify GFP-tagged protein from infected HaCat cells according to the manufacturer’s instructions. Briefly, infected cells were harvested at 24 hpi, washed in D-PBS then lysed in RIPA buffer. Cell lysates were centrifuged to clear cellular debris, before mixing with dilution buffer and adding GFP-TRAP beads. At this point a modification was made to the protocol: bead binding was performed overnight rather than for 1 h. Finally, the beads were washed three times before being resuspended in 2x SDS-PAGE lysis buffer and boiled for 5 minutes prior to analysis by SDS-PAGE.

### Quantitative RT-PCR (qRT-PCR)

Total RNA was extracted from cells using the RNeasy plus mini kit (Qiagen). Excess DNA was removed by incubation with DNase I (Invitrogen) for 15 min at room temperature, followed by inactivation for 10 min at 65°C in 25 nM of EDTA. Superscript III (Invitrogen) was used to synthesise cDNA using random primers according to the manufacturer’s instructions. All qRT-PCR assays were carried out in 96-well plates using Takyon™ No ROX SYBR 2X MasterMix blue dTTP (Eurogentec). Primers for cellular and viral genes are shown in S1 table. Cycling was carried out in a Lightcycler (Roche), and relative expression was determined using the ΔΔCT method (30), using 18s RNA as reference. Statistical analyses were carried out using unpaired t tests in GraphPad Prism 8.2.1.

### Immunofluorescence

Cells for immunofluorescence were grown on coverslips and fixed with 4%paraformaldehyde in PBS for 20 min at room temperature, followed by permeabilisation with 0.5% Triton-X100 for 10 min. Fixed cells were blocked by incubation in PBS with 10% newborn calf serum (block buffer) for 20 min, before the addition of primary antibody in block buffer, and a further 30-min incubation. After extensive washing with PBS, the appropriate Alexafluor conjugated secondary antibody was added in block buffer and incubated for a further 30 min. The coverslips were washed extensively in PBS and mounted in Mowiol containing DAPI. Images were acquired using a Nikon A1 confocal microscope and processed using ImageJ software (45).

### Fluorescent *in situ* hybridisation (FISH) of mRNA

Cells were grown in 2-well slide chambers (Fisher Scientific) and infected with virus. At the appropriate time, cells were fixed for 20 min in 4% PFA, then dehydrated by sequential 5 min incubations in 50%, 70% and 100% ethanol. FISH was then carried out using Applied Cell Diagnostics (ACD) RNAscope reagents according to manufacturer’s instructions. Briefly, cells were rehydrated by sequential 2 min incubations in 70%, 50% ethanol and PBS, and treated for 30 min at 37 °C with DNase, followed by 15 min at room temperature with protease. Cells were then incubated for 2 h at 40 °C with RNAscope probes for virus transcripts ICP27, TK and glycoprotein D, or cellular transcripts MMP1 and serpinE1, as designed by Advanced Cell Dignostics, ACD, followed by washes and amplification stages according to instructions. After incubation with the final fluorescent probe, the cells were mounted in Mowiol containing DAPI to stain nuclei, and images acquired with a Nikon A2 inverted confocal microscope and processed using Adobe Photoshop software.

**Table 1.**
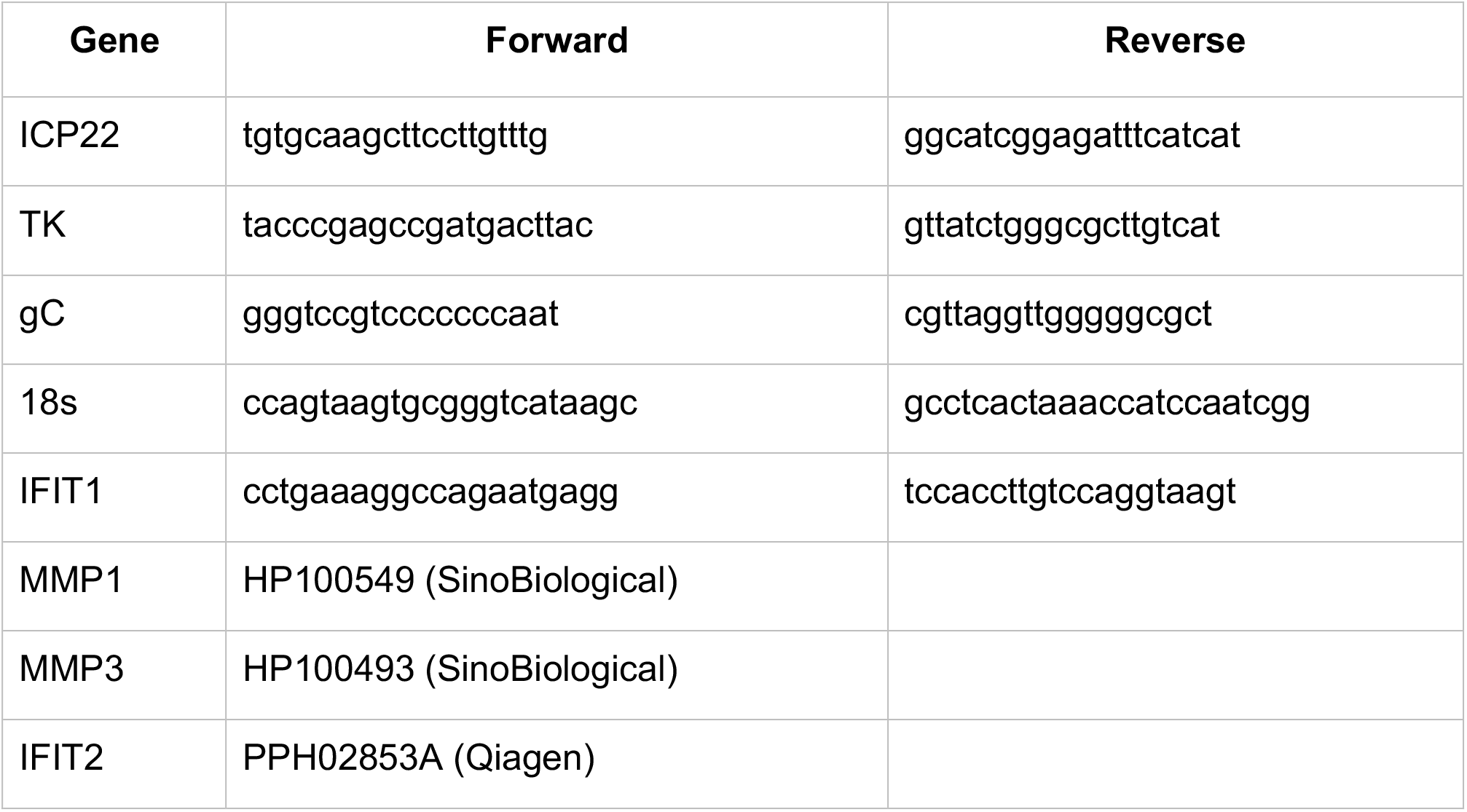
Primer pairs used for qRT-PCR (sequences shown 5’-3’)

## Acknowledgements

The authors would like to thank Colin Crump, Duncan Wilson and Frazer Rixon for antibodies used in this study. This work was funded by a grant from Medical Research Council, UK (MR/T0001038/1).

